# Predicting substitutions to modulate disorder and stability in coiled-coils

**DOI:** 10.1101/2020.06.04.131581

**Authors:** Yasaman Karami, Paul Saighi, Rémy Vanderhaegen, Denis Gerlier, Sonia Longhi, Elodie Laine, Alessandra Carbone

## Abstract

Coiled-coils are described as stable structural motifs, where two or more helices wind around each other. However, coiled-coils are associated with local mobility and intrinsic disorder. Intrinsically disordered regions (IDRs) in proteins are characterized by lack of stable secondary and tertiary structure under physiological conditions *in vitro*. They are increasingly recognized as important for protein function. However, characterizing their behaviour in solution and determining precisely the extent of disorder of a protein region remains challenging, both experimentally and computationally. In this work, we propose a computational framework to quantify the extent of disorder within a coiled-coil in solution and to help design substitutions modulating such disorder. Our method relies on the analysis of conformational ensembles generated by relatively short all-atom Molecular Dynamics (MD) simulations. We apply it to the phosphoprotein multimerisation domains (PMD) of Measles virus (MeV) and Nipah virus (NiV), both forming tetrameric left-handed coiled-coils. We show that our method can help quantify the extent of disorder of the C-terminus region of MeV and NiV PMDs, without requiring the input MD trajectory to actually sample the unfolded states of these regions. Moreover, this study provided a conceptual framework for the rational design of substitutions aimed at modulating the stability of the coiled-coils. By assessing the impact of four substitutions known to destabilize coiled-coils, we derive a set of rules to control MeV PMD structural stability and cohesiveness. We therefore design two contrasting substitutions, one increasing the stability of the tetramer and the other increasing its flexibility. Consequently, our method can be considered as a platform to reason about how to design substitutions aimed at regulating flexibility and stability.

## Background

Coiled-coils are ubiquitous oligomerisation motifs in proteins, where two or more amphiphatic *α*-helices intertwine together similarly to the strings of a rope. They account for approximately 5-10% of all protein-encoding sequences across all genomes [1]. The most common coiled-coils are left-handed and they feature a specific sequence motif called heptad repeat, consisting of seven residues *abcdefg* where *a* and *d* are hydrophobic [2]. The number of residues *per* turn in a regular *α*-helix is 3.6. In the case of heptad repeats, it reduces to 3.5 residues *per* turn, which leads to the left-handing [3]. A few cases of naturally occurring right-handed coiled coils were also identified, characterized by 11-residue repeats, where the periodicity of residues *per* turn increases up to 3.67. In this work we study the phosphoprotein multimerisation domains (PMD) of Measles virus (MeV) and Nipah virus (NiV) [4, 5], which both form tetrameric left-handed coiled-coils, and we compare their dynamical behaviour to that of the right-handed tetrameric coiled-coil of the RhcC protein (*Staphylothermus marinus*) [6].

MeV is a negative single stranded, non-segmented RNA virus that belongs to the family of *Paramyxoviridae* within the *Mononegavirales* order. Like in all *Mononegavirales* members, the genome of MeV is encapsidated in a regular array made of monomers of nucleoprotein (N) [7]. The N:RNA complex is the template for both transcription and replication ensured by the viral polymerase complex. The polymerase complex consists of the RNA-dependent RNA polymerase (L) and the phosphoprotein (P). P is an essential and conserved component of all non-segmented negative-sense RNA viruses, including some major human pathogens (e.g., rabies virus, respiratory syncytial virus, Ebola virus, and Nipah virus) [8]. All P proteins are multimeric, but the role of this multimerization is still unclear, with recent studies having reported that it is dispensable in the case of vesicular stomatitis virus [9, 10], a member of the *Rhabdoviridae family* in the *Mononegavirales* order, while PMD is crucially required for MeV polymerase functions [11]. MeV P is a modular protein [12], comprising an intrinsically disordered N-terminus domain (PNT, res 1-230) [13], and a C-terminal region (PCT, res 231-507). PCT is composed of a disordered region (res 231-303), the P multimerization domain (PMD) (res 304-375), a disordered linker (res 377-458) and a globular region (res 459-507) known as the X domain (XD) [14]. In the *wild-type* form of MeV PMD, residues in *a* and *d* positions are always leucine (L), isoleucine (I) and valine (V), except for N329 and Q356 (**Figure 1a**). The side chains of residues at these positions (*i.e. a* and *d*) point toward the interior of the coiled-coil and adopt a “knob into holes” packing, where a residue from one helix is accommodated into a hydrophobic pocket composed by the same residue from the three other chains (**Figure 1b**). The stability of coiled-coils is closely related to the geometry of knobs into holes. In MeV PMD, the regularity in the repetition of the *abcdefg* motif of the coiled-coil is interrupted by the insertion of a “stammer”, *i.e*. a three amino acids motif (L_339_L_340_L_341_) between two heptads. This stammer results in the formation of a short 3_10 helix that causes the coiled-coil to kink and exposes K343 (*e* position) to the solvent. Using truncation variants and biochemical assays, the C-terminal region of MeV PMD was found to be crucial for the chaperon activity of P towards the polymerase [11]. Those studies also showed that MeV gene expression relies on the cohesiveness of PMD coiled-coil, *i.e*. substitutions of hydrophobic *a* and *d* positions that modulate the stability of the PMD were found to affect viral gene expression [11].

**Figure 1:**
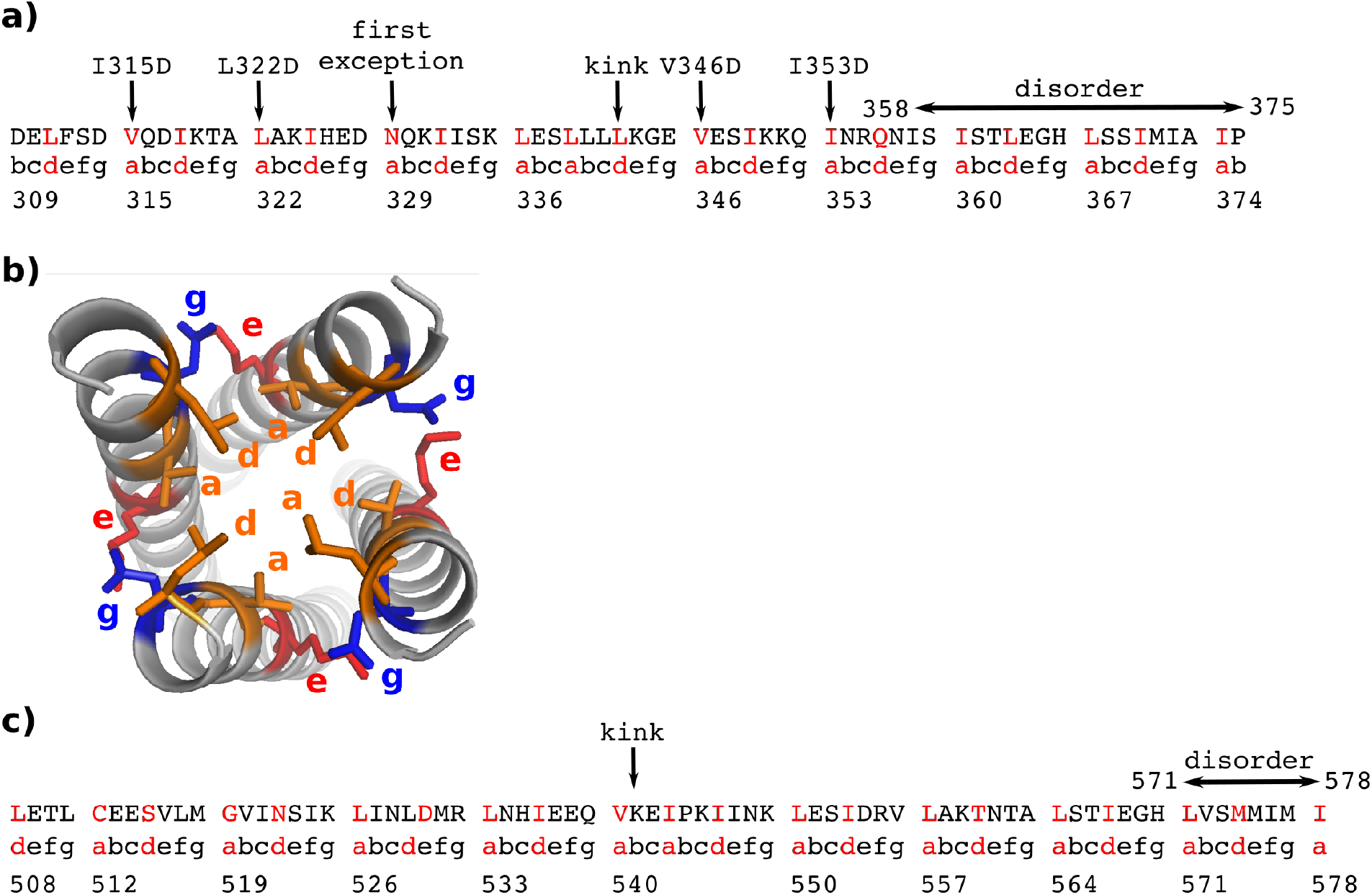
Sequence of registers of MeV PMD. The sequence of **a)** MeV PMD and **c)** NiV PMD along with the registers assigned to it are listed. Arrows point to the kink, the *a* positions of MeV PMD that have been substituted in this study, and the MeV PMD exception at position 329. **b)** The orientation of side chains are shown on the structure of the MeV PMD tetramer, for the hydrophobic (*a* and *d* registers) and charged (*e* and *g* registers) amino acids. Colors represent physico-chemical properties of amino acids: orange for hydrophobic, red for negatively charged and blue for positively charged.

The body of structural data available for MeV PMD are conflicting. The crystal structure of this domain has been solved by two groups. In one of these crystal structures, the full-length domain (residues 304-375) is well-ordered [15] (PDB code: 3ZDO), while in two others the C-terminal region (residues 358-375) is missing, consistent with structural disorder [4] (PDB code: 4C5Q and 4BHV). The details of the crystal structures are described in **Table I**. Structural comparison among the three available MeV PMD structures (4C5Q, 4BHV and 3ZDO) revealed crucial differences regarding the kink and the geometry of knobs [4]: *i*) the kink occurs at L342 in all chains of 4C5Q and 4BHV, whereas it is missing in chains C and F of 3ZDO, *ii*) the association of the tetramer is less tight and helices are significantly less twisted in 4BHV compared to the other two structures, *iii*) the structure of 3ZDO represents fewer knobs on each protomer and *iv*) all tetramers are energetically stable and the crystal structure of 4BHV has the highest stability. These findings illustrate how the same protein sequence may lead to different coiled-coil structures (*ie*. different content of disorder and different packing of the tetramer). Where do these differences come from? Are they due to different crystal packings, or to different conformations that PMD can adopt that might reflect different states within the L-P complex? Such differences can have an impact on the function and mechanism of action of this pivotal protein. It is therefore important to attempt at making sense of the discrepancies between the different structural data sets.

**Table I:** Structural details of the crystal structures of MeV and NiV PMDs. Three and two crystal structures are available for the PMD of MeV and NiV, respectively. Their structural details are reported.

| protein | PDB code | method | resolution<br>(Å) | chains | res. present<br>in the construct | res. resolved<br>in the structure |
| --- | --- | --- | --- | --- | --- | --- |
| MeV PMD | 3ZDO [15] | X-ray | 2.07 | A/B/C/D/<br>E/F/G/H | 304-377 | 308-371 |
|  | 4BHV [4] | X-ray | 2.10 | A/B/C/D/<br>E/F/G/H | 304-360 | 307-360 |
|  | 4C5Q [4] | X-ray | 2.20 | A/B/C/D | 304-375 | 307-357 |
| NiV PMD | 4N5B [5] | X-ray | 2.20 | A/B/C/D/<br>E/F/G/H | 470-578 | 475-578 |
|  | 4GJW [16] | X-ray | 3.00 | A/B/D/H/<br>E/F/G/H | 471-580 | 476-571 |

NiV is a newly emerged severe human pathogen in the family of *Paramyxoviridae* [17] and no vaccine or antiviral therapeutics has been developed yet for human use [18]. The N-terminal region of NiV P protein contains an extremely large intrinsically disordered region (residues 1-474) [19, 16]. Like the MeV P protein, the C-terminus region of NiV P encompasses a well-ordered multimerization domain (PMD), that spans residues 470-578, a flexible linker and a folded X domain (residues 660-709) [20]. The crystal structure of NiV PMD was solved as a long parallel tetrameric coiled-coil by two independent groups [5, 16]. In one of these crystal structures the coiled-coil region (residues 508-578) defined in the electron density is longer [5] (PDB code: 4N5B), while in the other the C-terminal region (residues 571-578) is missing [16] (PDB code: 4GJW), consistent with a disordered state like for MeV PMD. The details of the crystal structures are described in **Table I**. In striking contrast with MeV PMD, in NiV PMD the N-terminal region of each monomer forms a two-helix cap. Like for MeV PMD, the regularity of the heptad repeat of the coiled-coil is interrupted by a stammer, *i.e*. the insertion of the I_543_P_544_K_545_ motif, and a kink occurs on residues V_540_K_541_E_542_ (**Figure 1c**).

Molecular dynamics (MD) simulations provide a powerful way to describe the dynamical behaviour of proteins in solution. However, the size of a protein’s conformational space grows exponentially with the number of atoms and its full exploration remains prohibitive, especially when the protein is intrinsically disordered [21]. We have previously developed a method, COMmunication MApping (COMMA) that facilitates the description of protein dynamical architecture starting from conformational ensembles generated by relatively short MD simulations [22]. A more recent version of the tool (COMMA2) further allows predicting substitution effects at large scale and identify positions highly sensitive to substitutions [23]. In the present work, we used COMMA2 to resolve the ambiguities in the crystal structure data, and to assess the effects of several substitutions on the structural stability of the tetramers. First, we show that COMMA2 is useful to detect protein regions prone to disorder without requiring the input MD trajectory to actually sample the unfolded states of these regions. Second, we demonstrate that substituting a set of *a* positions with negatively charged residues modulates the structural cohesiveness of the MeV PMD tetramer in a position-specific manner. These results provide an asset to rationally design two new substitutions modulating the stability of the coiled-coil. Specifically, targeting the kink increases its flexibility while targeting the C-terminus increases its stability.

## Methods

### Proteins studied

We studied three homo-tetrameric coiled-coils whose structures were resolved by X-ray crystallography: *i)* MeV PMD (PDB id: 3ZDO, residues 308-373, 2.07Å resolution), *ii)* NiV PMD (PDB id: 4N5B, residues 475-578, 2.2Å resolution) and RhcC (PDB id: 1YBK, residues 1-52, 1.45Å resolution). Among the available crystal structures for MeV PMD and NiV PMD, we chose 3ZDO and 4N5B, respectively, since they both have the largest number of residues defined in the electron density and with a C-terminus that is resolved. The right-handed coiled-coil homo-tetramer of the RhcC protein was taken as a control for our analysis on left-handed coiled-coils. Furthermore, we studied 6 different substitutions of MeV PMD: V315D, L322D, L336D, V346D, I353D, and E364F, hence in total 9 systems.

### Molecular dynamics simulations

#### Set up of the systems

The 3D coordinates for the studied proteins were retrieved from the Protein Data Bank (PDB) [24]. All crystallographic water molecules and other non-protein molecules were removed. All models were prepared using the LEAP module of AMBER 12 [25], with the ff12SB forcefield parameter set: (*i*) hydrogen atoms were added, (*ii*) Na^+^ or Cl^-^ counter-ions were added to neutralise the system charge, (*iii*) the solute was hydrated with a cuboid box of explicit TIP3P water molecules with a buffering distance up to 10Å. The environment of the histidines was manually checked and they were consequently protonated with a hydrogen at the *ϵ* nitrogen. The variants of MeV PMD were generated by *in silico* substitutions starting from the 3ZDO structure using Rosetta Backrub [26].

#### Minimisation, heating and equilibration

The systems were minimised, thermalised and equilibrated using the SANDER module of AMBER 12. The following minimisation procedure was applied: (*i*) 10,000 steps of minimisation of the water molecules keeping protein atoms fixed, (*ii*) 10,000 steps of minimisation keeping only protein backbone fixed to allow protein side chains to relax, (*iii*) 10,000 steps of minimisation without any constraint on the system. Heating of the system to the target temperature of 310 K was performed at constant volume using the Berendsen thermostat [27] and while restraining the solute *C_α_* atoms with a force constant of 10 kcal/mol/Å^2^. Thereafter, the system was equilibrated for 100 ps at constant volume (NVT) and for further 100 ps using a Langevin piston (NPT) [28] to maintain the pressure. Finally the restraints were removed and the system was equilibrated for a final 100-ps run.

#### Production of the trajectories

For every protein, 2 replicates of 50 ns, with different initial velocities, were performed in the NPT ensemble using the PMEMD module of AMBER 12. The temperature was kept at 310 K and pressure at 1 bar using the Langevin piston coupling algorithm. The SHAKE algorithm was used to freeze bonds involving hydrogen atoms, allowing for an integration time step of 2.0 fs. The Particle Mesh Ewald method (PME) [29] was employed to treat long-range electrostatics. The coordinates of the system were written every ps. Standard analyses of the MD trajectories were performed with the *ptraj* module of AMBER 12.

#### Stability of the trajectories

The root mean square deviations (RMSD) of the studied coiled-coils (MeV PMD wild-type and variants, NiV PMD and RhcC) were measured along simulation time for all the replicates (**Figures S1** and **S2**). All systems were fully relaxed after 10 ns. Consequently, the last 40 ns of each replicate were retained for subsequent analyses. Moreover, we calculated the residue root mean square fluctuations (RMSF) over the last 40 ns of every simulation (**Figure S3**).

### COMMA2 analysis

For every studied system, COMMA2 was applied to the last 40 ns of the two replicates of MD simulations, and communication blocks were extracted. COMMA2 identifies pathway-based communication blocks (*CBs^path^), i.e*. groups of residues that move together, and are linked by non-covalent interactions, and clique-based communication blocks (*CBs^clique^*), *i.e*. groups of residues that display high concerted atomic fluctuations, and that are close in 3D space (see [22] for formal definitions and detailed descriptions). It should be emphasized that all the backbone-backbone non-covalent interactions are ignored for this analysis. We defined two sets of *CBs^path^*, namely short-range and long-range blocks [22]. Short-range blocks consist of pathways of at least 4 residues. Long-range blocks are system-dependent, *i.e*. they consist of pathways with at least 7 or 8 residues. COMMA2 detects pairs of stable secondary structure elements directly linked by communication pathways. Moreover, we modified the definition of thresholds, and implemented new metrics and functionalities that allowed us to identify disordered regions, as explained below.

#### Identification of communication pathways

*Communication pathways* are chains of residues that are not adjacent in the sequence, and form stable non-covalent interactions (hydrogen-bonds or hydrophobic contacts), and communicate efficiently. *Communication efficiency or propensity* is expressed as [22]:

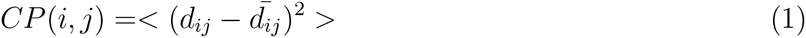

where *d_ij_* is the distance between the *Cα* atoms of residues *i* and *j* and 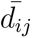 is the mean value computed over the set of conformations. Two residues *i* and *j* are considered to communicate efficiently if *CP*(*i,j*) is below a *communication propensity threshold*, CP_*cut*_. The strategy employed to set the value of *CP_cut_*, is explained in [22]. However, the algorithm is modified in this work by considering the definition of chains. Intuitively, it is expected that neighbouring residues in the sequence, that form well-defined secondary structures, communicate efficiently with each other. Therefore, we evaluate the proportion *p_ss_* of residues that are in an *α*-helix, a *β*-sheet or a turn in more than half of the conformations. Then for every residue *i* surrounded by 8 sequence neighbours (4 before and 4 after), we compute a *modified communication propensity MCP*(*i*) as:

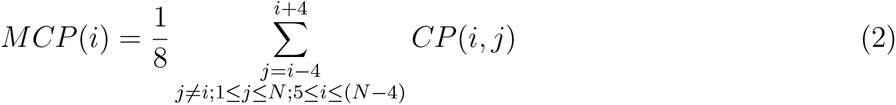

where *N* is the total number of residues in each chain. *CP_cut_* is chosen such that the proportion *p_ss_* of *MCP* values are lower than *CP_cut_*. Whenever more than one replicate of MD trajectories are available, we measured the *CP_cut_* for each replicate and considered the average value for the identification of pathways. In addition, for the variants of MeV, we used the same *CP_cut_* as the wild type to better contrast their behaviours.

#### Residue confidence scores

*Communication pathways* and *independent cliques* are defined by setting several parameters [22]. To assess the robustness of the results obtained with the default parameter values, COMMA2 systematically explores the parameter space around those values and assigns *confidence scores* to every residue in a *CB^path^* or *CB^clique^* [23]. The default procedure is to vary the thresholds from their default value up to the value where all residues of the protein are in the same block.

#### Prediction of disorder using COMMA2

A *CB^clique^* is considered as predictor of disorder, if the residues forming that block encompass several chains. To reflect this behaviour, we propose two scores: residue disorder propensity (*RDP*) and disorder clique score (*DCS*). *RDP* highlights the propensity of each residue *j* for belonging to *CBs^clique^* encompassing several chains, and is measured as:

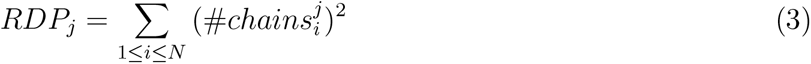

where N is the number of *CBs^clique^* and 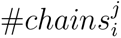 is the number of chains in 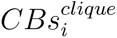 at position *j*. Hence, the *N*^2^ corresponds to the highest disorder propensity. *DCS* accounts for the disorder propensity of each 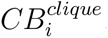, and is calculated as:

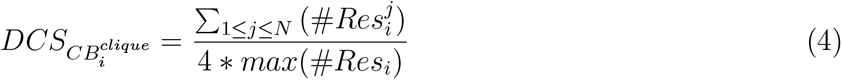

where N is the number of chains and 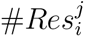 is the number of residues in 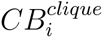 and in chain *j*. If the numbers of residues from each chain are very different, the score is very small, and if the same number of residues from each chain are present in a *CB^clique^*, the score equals to one, highlighting the disorder region.

### Comparison with other approaches

We compared the performance of our method with four other existing approaches to predict either coiled-coils or disordered regions in proteins, namely Coils server [30], IUPred [31], DISOPRED [32] and Rosetta ResidueDisorder [33]. The Coils server measures the probability of a sequence to adopt a coiled-coil conformations. It takes a sequence as input and compares it with a database of known parallel two-stranded coiled-coils and measures a score based on the similarity. Consequently, it compares the score with the distribution of scores for globular and coiled-coils proteins to obtain the probability of forming coiled-coil conformation. IUPred predicts intrinsically disordered regions based on estimation of the pairwise energy content. Amino acids in globular proteins are able to form a large number of favorable interactions, while intrinsically disordered proteins do not have the potential to form enough favorable interactions, due to their lack of stable structure. DISOPRED predicts disordered regions based on disorder data from high resolution X-ray crystal structures, *i.e*. residues missing in the electron density map, but present in the sequence record. DISOPRED exploits support vector machine (SVM) learning techniques to train a neural network classifier and estimates the probability of residues being disordered. Finally, Rosetta ResidueDisorder uses Rosetta to predict the structure of a protein and reports the Rosetta per-residue energy function scores. Intrinsically disordered regions have higher score than ordered regions.

## Results and discussion

For each system of MeV PMD, NiV PMD and RhcC, we performed two replicates of 50 ns MD simulations, and applied COMMA2 to extract communication blocks. COMMA2 allows to contrast the different types of communication occurring between residues and to hierarchize the different regions of a protein depending on their communication efficiency. To this end, COMMA2 identifies pathway-based communication blocks (*CBs^path^*) and clique-based communication blocks (*CBs^clique^)*. The residues comprised in a *CB^path^* are linked by *communication pathways* by transitivity. A communication pathway is defined as a chain of residues displaying correlated motions and linked by stable non-covalent interactions (see *Methods* and [22] for more details). Hence, it represents an efficient route of information transmission supported by physical interactions. By contrast, *CBs^clique^* are the most flexible regions of the protein displaying highly concerted atomic fluctuations. The RhcC protein was chosen as a control for our analysis on left-handed coiled-coils, for two reasons: *i*) the right-handed nature of this homo-tetrameric coiled-coil and *ii*) the lack of kink in the middle of the structure.

### Identical chains differentially contribute to the stability of MeV and NiV PMD tetramers

COMMA2 analysis revealed that, although the sequence of amino acids for each studied tetramer, is identical between its four chains, the behaviour of the chains within MeV and NiV PMDs is different. In the case of MeV PMD, we observe two *CBs^path^* (**Figure 2a**), of which one (in red) contains most of the residues from the four chains. A similar behaviour is observed for NiV PMD (**Figure 2d**). By contrast, for RhcC only one *CB^path^* is observed, suggesting a symmetrical/equivalent role for the different chains (**Figure 2g**). The hierarchy of communication pathways between the helices in MeV and NiV PMD, can be further refined by considering only long pathways (**Figure 2b,e**). We can clearly see that long-range communication occurs only in the N-terminal half of the tetramers. In contrast, the long-range *CB^path^* of RhcC encompasses residues from the 4 chains and both halves (71% of the total number of residues in the protein).

**Figure 2:**
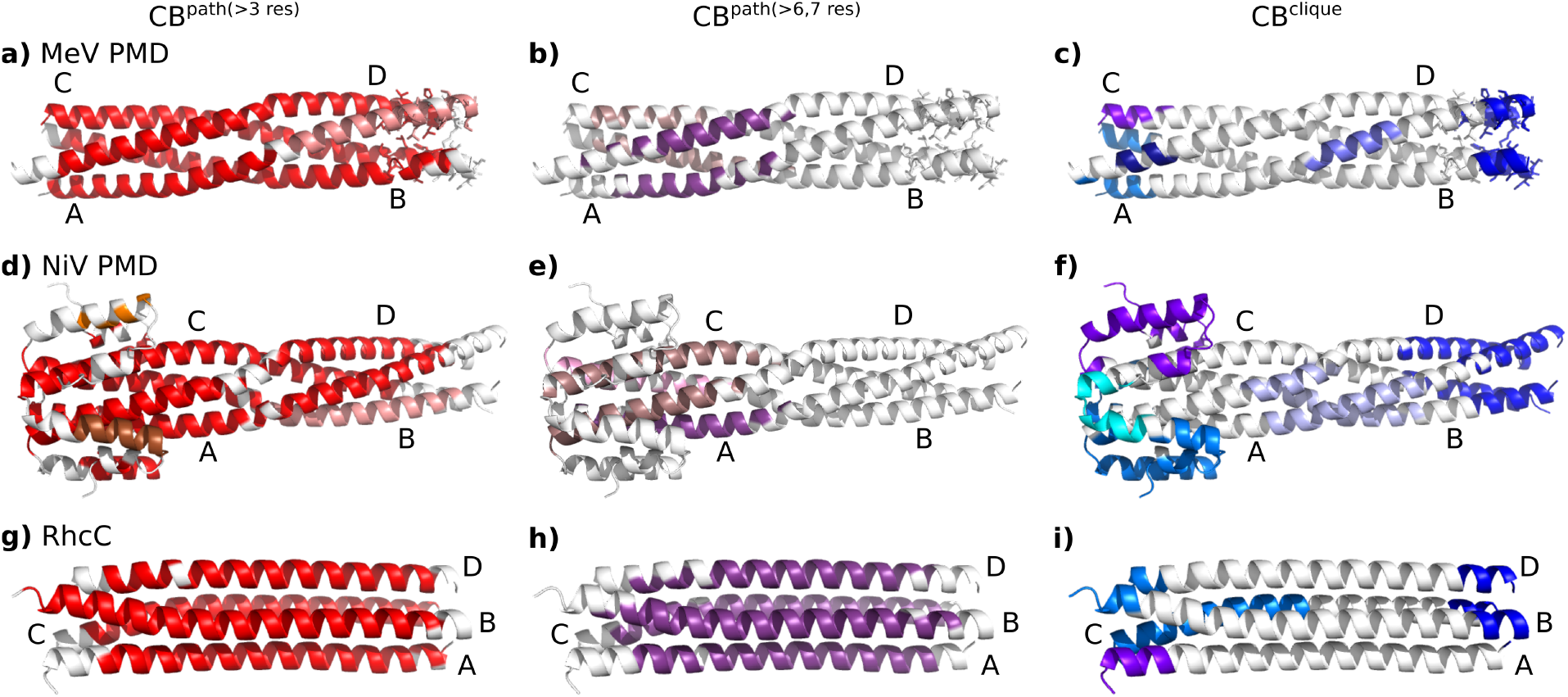
Communication Blocks of MeV PMD, NiV PMD and RhcC identified by COMMA2. *CBs^path^* and *CBs^clique^* identified by COMMA2 are mapped on the structure of (**a,b,c**) MeV PMD (PDB code: 3ZDO), (**d,e,f**) NiV PMD (PDB code: 4N5B) and (**g,h,i**) RhcC (PDB code: 1YBK). Left (**a,d,g**): distinct *CBs^path^* of at least 4 residues are colored in shades of red. Middle (**b,e,h**): long-range *CBs^path^* are depicted in shades of purple, containing at least 7 residues (for NiV PMD) or 8 residues (for MeV PMD and RhcC). Right (**c,f,i**): distinct *CBs^clique^* are colored in blue tones. Residues known to be ambiguous in MeV PMD are shown by sticks.

Regarding *CBs^clique^*, they are detected in three locations in MeV and NiV PMDs: at the N-term, around the kink (kink-region) and at the C-term. We observed a striking contrast between the N- and C-termini. While single-chain CBs^*clique*^ are detected in the former, only one *CB^clique^* encompassing residues from all four helices is detected in the latter (**Figure 2c,f**, in dark blue). In the case of MeV PMD, 65% of the residues known to be ambiguous, *i.e*. missing in one crystal structure (shown as sticks on **Figure 2c**) are included in this block. Hence, the detection of this *CB^clique^*, indicative of concerted high fluctuations across the four chains, can be interpreted as an indicator of disorder. We can thus hypothesize that residues of the corresponding block in NiV PMD are, to some extent, disordered. These results are in agreement with the existence of an ambiguous region with higher flexibility in the C-terminal end of NiV, that was discussed in a recent study [5]. All those structural ambiguous residues are included in the C-term *CB^clique^* of NiV PMD. In contrast, COMMA2 results of the right-handed RhcC suggest a strongly different behaviour from what we observed for the left-handed coiled-coils (**Figure 2i**). The four N-termini of the tetramer move independently from each other since none of the identified *CB^clique^*s at this region, encompasses the four helices. In addition, the *CB^clique^* at the C-terminus, contains only one residue from chain A. This observation suggests the lack of a disordered region in the structure of RhcC. The observed contrast between MeV and NiV PMDs and our negative control RhcC, highlights the impact of the kink in the middle of the left-handed tetramers.

To investigate whether differences could also be detected in a *monomeric context*, we performed MD simulations on single chains extracted from the tetramer of MeV PMD, NiV PMD and RhcC, and applied COMMA2 on each system (details are reported in **Supporting Materials**). We should specifically stress that we did not intend here to realistically simulate the behaviour of a single helix in solution. As expected, the single helices were very flexible in the simulations, reflecting their instability. Nevertheless, we observed some differences suggesting that MeV and NiV PMD “monomers” were more prone to unfolding at their C-terminus than RhcC “monomer” (**Figure S4**).

### COMMA2 as a predictor of disorder and comparison with other tools

Based on the previous results, we propose a formal criterion to predict regions prone to be disordered in otherwise folded coiled-coils. We define a residue disorder propensity (*RDC*) index reflecting the propensity of each residue to be part of a *CB^clique^* encompassing several chains (**Figure 3**, *cf* regions in blue with the experimentally detected ambiguous regions, in black). In order to evaluate the power of COMMA2 to predict disordered region in coiled-coil structures, we compared our results with random coil propensities estimated by DSSP [34] (in red) and with predictions from four software: Coils server, IUPred, DISOPRED and Rosetta ResidueDisorder. The Coils server measures the probability to form a coiled-coil structure. Here we chose to present the probability not to form a coiled-coil structure (which is 1 – *Prob*(*CoilsServer*)), in order to better compare the results (in green). IUPred and DISOPRED both predict the probability of residues to be disordered (orange and purple bars, respectively). Rosetta ResidueDisorder predicts ordered (in white) and disordered residues (in magenta).

**Figure 3:**
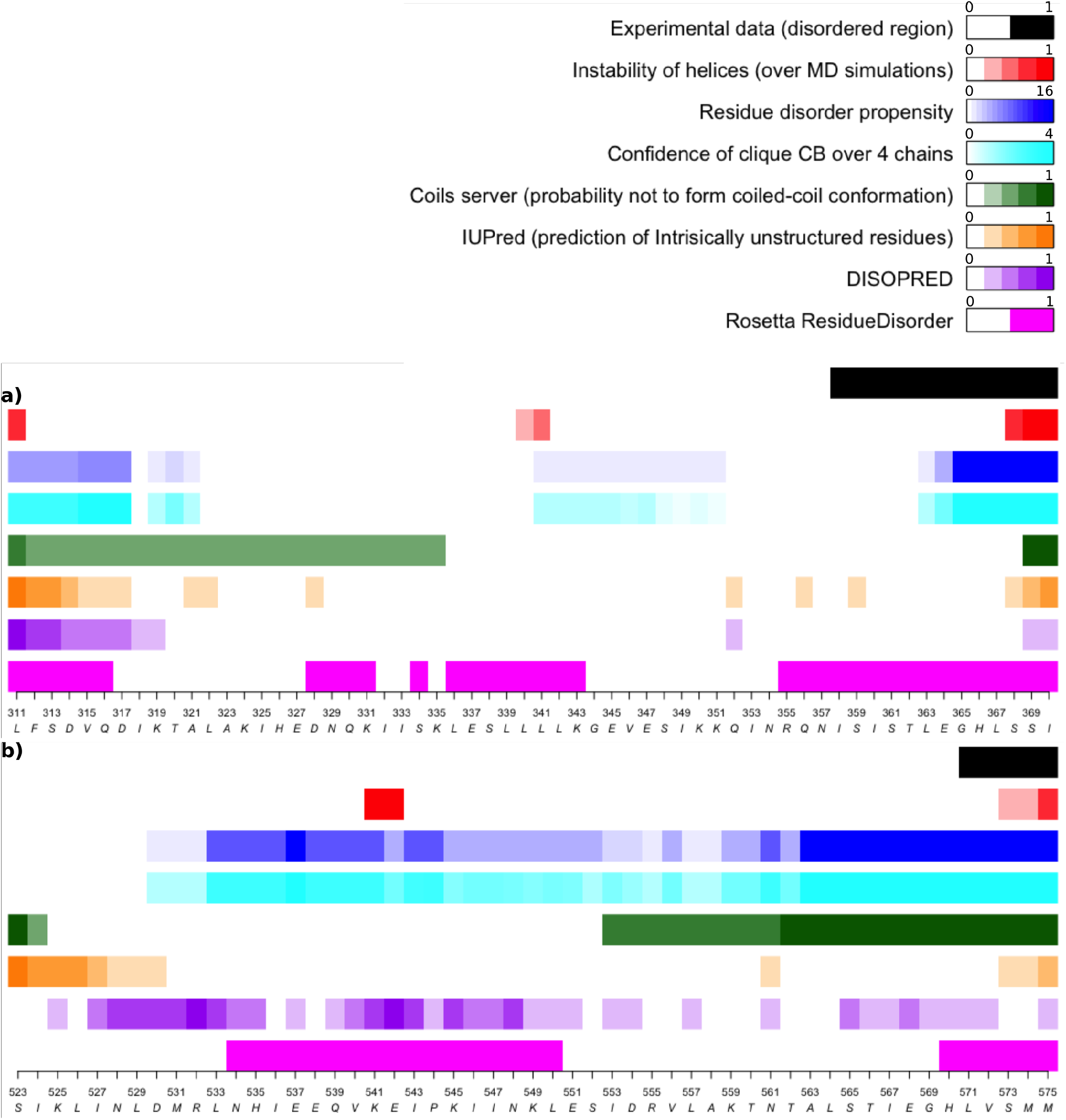
Comparison between different methods predicting disordered residues. Comparison between different methods predicting disorder residues for **a)** MeV PMD and **b)** NiV PMD. The unfolding of helices along MD simulations, the propensity of residues detected as *CB^clique^* and the confidence of *CB^clique^* residues are shown in red, blue and cyan, respectively. The predicted propensity of the Coils server to form coiled-coil structures is shown in green. The predictions of disordered residues obtained from IUPred, DISOPRED and Rosetta ResidueDisorder are shown in orange, purple and magenta, respectively.

In the case of MeV PMD, the comparison of the predictions with the experimental data, reveals the power of COMMA2 to predict the disordered residues with strikingly better accuracy compared to Coils server, IUPred and DISOPRED (an accuracy of 62% compared to 15%, 23% and 0%, respectively). Rosetta ResidueDisorder displays the highest sensitivity, but it also detects other regions of the coiled-coil as disordered, for instance residues at the N-terminus or in the kink-region. In the case of NiV PMD, the accuracy of COMMA2, Coils server and Rosetta ResidueDisorder are all 100%, while IUPred and DISOPRED both fail to predict the C-terminus residues as being prone to disorder (**Figure 3b**).

While the sequence-based methods (Coils and IUPred) and DISOPRED are good and fast measures to predict coiled-coil (Coils server) and disorder (IUPred and DISOPRED) propensities, they are not able to detect unstable regions of coiled-coils. On the other hand, Rosetta ResidueDisorder has high sensitivity but low positive predictive value (PPV) to predict disordered residues, while being computationally slower compared to the other three methods. Although COMMA2 is computationally more expensive as it relies on the results of MD simulations, it is able to predict both flexible and unstable regions. Interestingly, COMMA2 also provides a mean to distinguish the two different behaviours: while the presence of a single *CB^clique^* over the four chains reflects the propensity to local disorder, the existence of *CBs^clique^* provides hints on flexible regions playing a role in coiled-coils dynamics.

### Controlling MeV PMD flexibility/communication through substitutions

The main purpose of this study is to design some substitutions modulating the stability of the structure of MeV PMD. The *a* positions on the sequence of MeV PMD are particularly attractive for this, as they bear hydrophobic side chains oriented toward the interior of the coiled-coils (inward). Consequently, substituting those positions with a negatively charged amino acid is expected to bring some instability by disrupting the “knobs into holes” organization that relies on hydrophobic interactions (**Figure 4)**. The analysis of systematic substitution of a coiled coil has shown that similar substitutions of hydrophobic *a* positions with negatively charged amino acids, destabilize the structure of coiled-coils [35], a feature that can be associated with function (see [36] for example). However, the structural effects that such substitutions would induce are not fully understood. Therefore, COMMA2 analysis can be employed to highlight the details of interaction networks and communications across the structure, unveiling structural changes induced by different sets of substitutions. From the analysis of the wild-type PMD structure, one may infer asymmetric dynamics of the protein, *i.e*. the first half of the structure is more rigid and represents the communication core of the system, whereas the second half is more flexible and contains disordered residues. Here, we investigated how substitutions creating local instability could influence the communication efficiency throughout the tetramer and in particular between the two halves. Specifically, we systematically substituted the hydrophobic amino acid at each *a* position (**Figure 4a**) with the negatively charged amino acid D and thus investigated the impact of the following substitutions: V315D and L322D that fall in the first half of the helices, and V346D and I353D in the second half of the helices (**Figure 4b**). Upon replacement of the wild-type residue with an aspartic residue, two replicates of 50 ns MD simulations were performed for every variant. COMMA2 analysis was applied to the ensemble of conformations obtained for each variant. The results suggest that substitutions in the first half (V315D and L322D), establish/reinforce communication between the two halves and the roles of the two halves in the communication of the complex becomes almost symmetrical/equivalent. In contrast, substitutions in the second half (V346D and I353D), result in the breakage of the communication between the two halves. Consequently, the first half becomes more rigid, while the flexibility of the second half increases. In the following the detailed results are reported.

**Figure 4:**
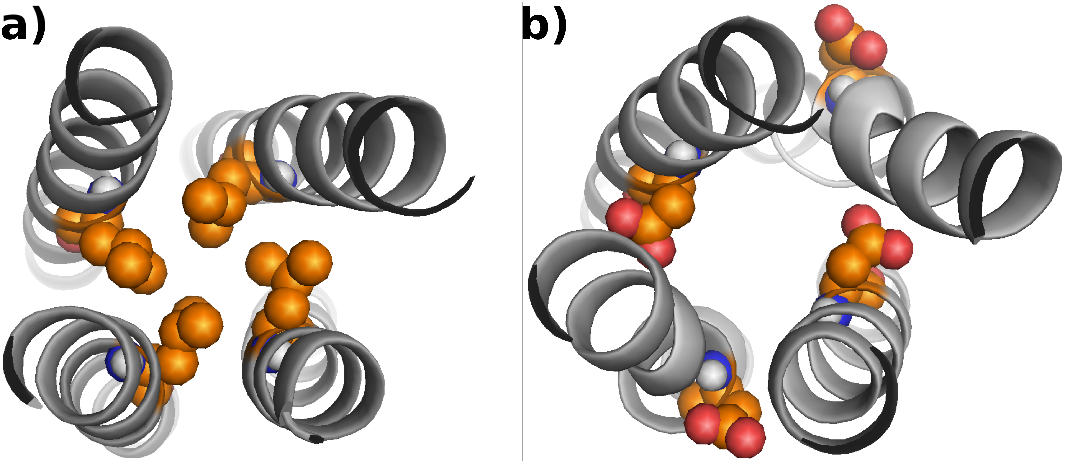
Substitution of a hydrophobic amino acid with a negatively charged one. The hydrophobic *a* positions (**a**) are substituted to negatively charged residues (**b**). Corresponding side chains are shown as spheres.

### Substitutions in the first half, before the kink

The V315D and L322D substitutions are located in the N-terminal extremity, positioned in the first and second *a* register, respectively. We applied COMMA2 analysis to extract the set of *CBs^path^* and *CBs^clique^*, to measure the direct communications between pairs of secondary structure elements (see *Methods*), and to compare the results with the wild-type PMD (**Figure 5)**. The main differences between the variants and the wild type are: (1) the residues preceding the substitution are excluded from the main *CB^path^* (in red), (2) the splitting of the helices into two groups, when considering long pathways, is not observed anymore and we detect only one long-range *CB^path^* (in purple), (3) the latter gets shifted toward the second half as the position targeted for substitution is shifted towards the C-terminal end, (4) this is accompanied by an increase in the number of direct communications between the two halves (**Figure 5e, f** and **Table II**), (5) the *CB^clique^* observed near the kink disappears. Moreover, we estimated the disorder tendency of the *CBs^clique^*, through the disorder clique score (*DCS*), which reflects the ability of a *CB^clique^* to encompass residues from several chains (see *Methods*). We observed that as the position targeted for substitution is shifted towards the C-terminal end, the *CB^clique^*, detected at the N-terminus become more indicative of disorder (**Figure 5d**). We can conclude that substituting the hydrophobic *a* positions on the first half with a negatively charged amino acid, establishes/reinforces communication between the two halves. The roles of the two halves in the communication of the complex become almost symmetrical/equivalent.

**Table II:** Summary of the number of residues involved in *CBs^path^, CBs^clique^* and in direct pathways. For the wild type and six variants of MeV PMD, the details of the *CBs^path^* and *CBs^clique^* are reported. In addition, the number of direct communications between the chains in the two halves is also shown.

| | $CBs^{path}$ | $CB^{clique}$ | | | direct pathways | |
| --- | --- | --- | --- | --- | --- | --- |
|  |  | N-term | 2nd half | C-term CB | N-term | C-term |
| wild type | 195, 25 | 19, 8, 6 | 11 | 32 | 35 | 11 |
| V315D | 209 | 28, 9 | - | 23 | 65 | 17 |
| L322D | 166, 24 | 25 | - | 32 | 58 | 31 |
| L336D | 95, 89 | 16 | - | 36 |  |  |
| V346D | 123, 20, 19, 19, 19 | 15, 9 | 19 | 30 | 63 | 4 |
| I353D | 169, 16, 4, 18 | 17 | 20 | 36 | 55 | 12 |
| E364F | 211 | 24 | - | 33 |  |  |

**Figure 5:**
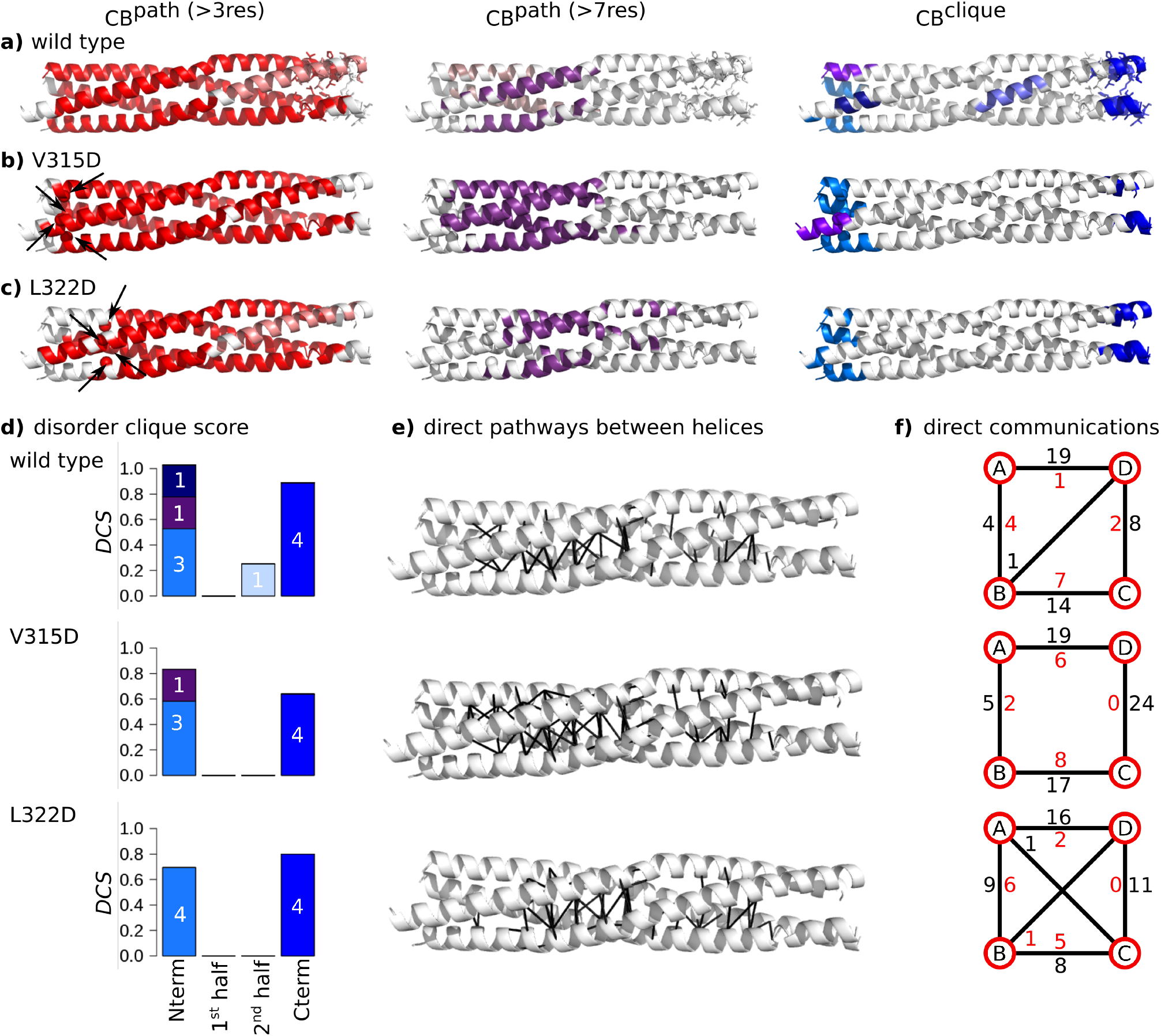
Inter-residue communication in V315D and L322D variants of MeV PMD. Communication blocks identified by COMMA2 are mapped on the average MD conformation for the wild type (**a**) and two variants bearing substitutions in the first half of the helices, V315D (**b**) and L322D (**c**). *CBs^path^* are obtained by considering interactions involving side chains and pathways with length equal or greater than 4 and 8 residues. Residues known to be disordered in a contextdependent manner are shown as sticks. The *CBs^clique^* identified by COMMA2 are colored in blue tones. **d)** The disorder clique score (*DCS*, see *Methods*) is reported for every *CBclique* of the wild type and variants of MeV PMD. **e)** All direct communications between helices are shown on the structures. **f)** The schematic representation, reports the number of direct pathways between helices in the first and second halves (numbers colored in black and red, respectively).

### Substitutions in the second half, after the kink

The V346D and I353D substitutions are located in the C-terminal extremity, with V346D being very close to the kink. Similar to the previous substitutions, we applied COMMA2 analysis to extract the sets of *CBs^path^* and *CBs^clique^*, and direct communications between pairs of secondary structure elements (**Figure 6)**. The main differences between the variants and the wild type are: (1) the biggest *CB^path^* (in red) is smaller than the corresponding block in the wild type and new blocks appear downstream the substitution position in the C-terminal half (in shades of pink), (2) considering long pathways, the splitting of the helices into two groups is not observed anymore, and we detect only one long-range *CB^path^* (in purple), (3) this is accompanied by the lack or presence of only few direct communications between the chains in the second half (**Figure 6e,f** and **Table II**), (4) the number and size of the *CBs^clique^* in the N-terminal end decrease, and (5) the *DCS* of *CB^clique^* observed near the kink increases as it spans residues from two chains (**Figure 6d**). We can conclude that substituting the hydrophobic *a* positions in the second half with a negatively charged amino acid, induces a breakage of communication between the first and second halves, thereby impending the propagation of communication across the structure. The structure is more rigid in the first half while it is even more flexible in the second half and chains in the second half exhibit a more independent behaviour.

**Figure 6:**
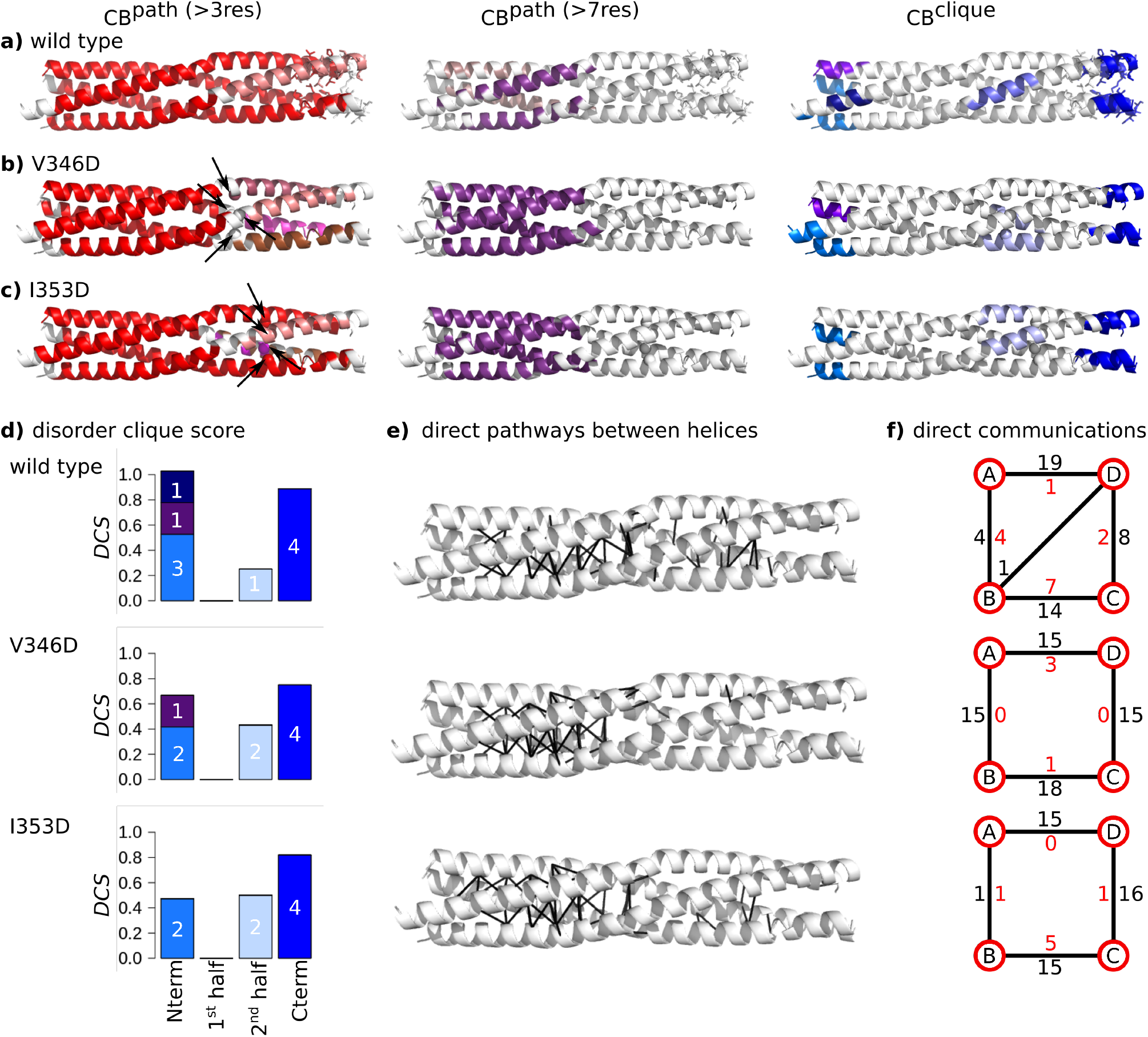
Inter-residue communication in V346D and I353D variants of MeV PMD. Communication blocks identified by COMMA2 are mapped on the average MD conformation for the wild type (**a**) and for variants bearing substitutions in the second half of the helices, I353D (**b**) and V346D (**c**). Those pathways are obtained by considering interactions involving side chains and pathways with length equal or greater than 4 and 8 residues. Residues known to be disordered in a context-dependent manner are shown by sticks. The *CBs^clique^* identified by COMMA2 are colored in blue tones. **d)** The disorder clique score (*DCS*) is reported for every *CBclique* of the wild type and variants of MeV PMD. **e)** All direct communications between helices are shown on the structures. **f)** The schematic representation, reports the number of direct pathways between helices in the first and second halves (numbers colored in black and red, respectively).

### Educated guess of substitutions modulating MeV and NiV PMD stability

COMMA2 analysis of the impact of various substitutions within MeV PMD elucidates the asymmetric dynamics of MeV PMD, and the role of different regions on the overall stability of the helices. Therefore, COMMA2 can be considered as a platform to reason about how to design amino-acid changes aimed at regulating flexibility and stability. Indeed, we are able to extract general rules for designing substitutions that modulate the overall stability: *i)* the first half of the tetramer is more rigid, therefore introducing additional perturbations (substituting the hydrophobic *a* positions) in this region should result in symmetrical behaviour between the two halves of the tetramer and increase of overall stability, *ii)* the kink-region plays a crucial role in the propagation of communications across the tetramer, therefore perturbations at this region may lead to the asymmetry and disruption of the communications, *iii)* the second half is more flexible and substituting the hydrophobic *a* positions in this region, should increase the asymmetric dynamics and the overall flexibility, and *iv)* in contrast, a reverse substitution, *i.e*. from a negatively charged amino acid to a hydrophobic one, should increase the symmetry between the two halves. In this section, based on these rules we propose two contrasting substitutions, one increasing the flexibility, and the other bringing more stability.

The presence of a stammer (L_339_L_340_L_341_) in MeV PMD induces the formation of a kink at these positions. Based on the results from the previous section, it appears that the kink plays a critical role in the communication between the two halves of the helices, a finding in agreement with the critical functional role that the kink plays in transcription and replication [11]. We thus reasoned that substituting hydrophobic positions close to the kink, may disrupt the communications. For this reason we designed a variant in which L336 was replaced with an aspartate (L336D). We applied COMMA2 and identified two *CBs^clique^* (**Figure 7a**). The C-term clique spans the four chains and contains a slightly increased number of residues (36 aa) compared to the wild type (**Figure 7c**). The other N-terminal clique encompasses residues from three chains (16 aa). Instead of displaying one large *CB^path^* spanning the two halves, like the wild type, this variant shows a sharp division between the two halves, with one *CB^path^* located exclusively in the first half and the other one exclusively in the second half. Considering long-range pathways (at least 8 residues), two small blocks are detected, with a different pairing of the chains. The analysis of direct communication between helices (**Figure 7d,e**) highlights a significant decrease of such communications in the first half, where the contacts are all concentrated in narrow/small regions (they are very localized and not spread over the length of each half). Overall, these results suggest an increase of flexibility at both halves and disruption of communication around the substituted position and the kink.

**Figure 7:**
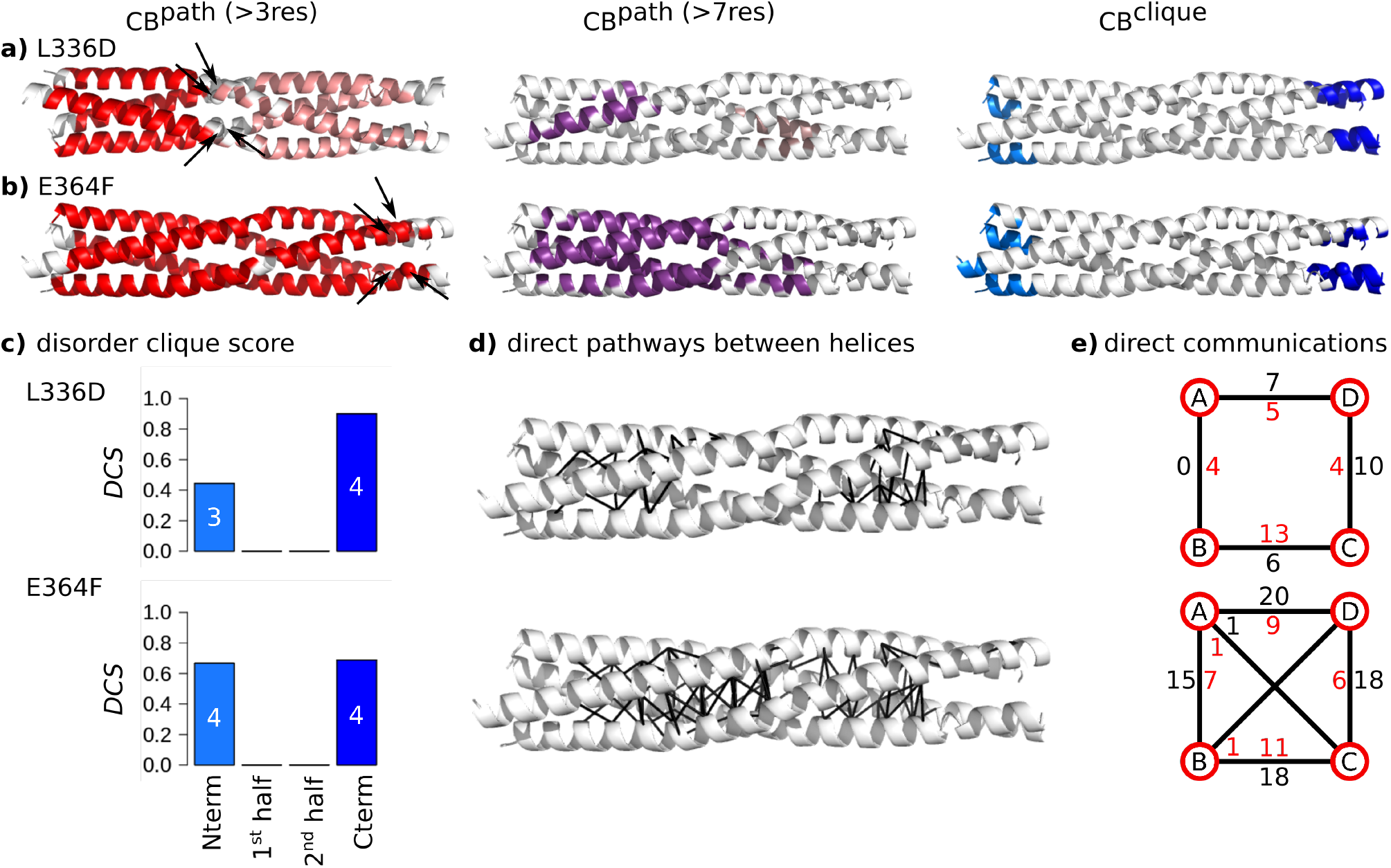
Inter-residue communications in L336D and E364F variants of MeV PMD communication. Communication blocks identified by COMMA2 for **a)** L336D and **b)** E364F MeV PMD variants. *CBs^path^* are obtained by considering interactions involving side chains and pathways with length equal or greater than 4 and 8 residues. The *CBs^clique^* identified by COMMA2 are colored in blue tones. **c)** The disorder clique score (*DCS*) is reported for every *CBclique*. **d)** All direct communications between helices are shown on the structures. **e)** The schematic representation, reports the number of direct pathways between helices in the first and second halves (numbers colored in black and red, respectively).

Next, we designed a substitution expected to bring more stability to the communication network of MeV PMD. To do this, we targeted a negatively charged residue from the disordered region (E364) and replaced it with a phenylalanine (a hydrophobic and bulk amino acid). We expected that this substitution would bring the side chains at position 364 at the interface of helices, thereby increasing the overall communication. COMMA2 analysis of E364F, revealed two *CBs^clique^* (**Figure 7b**). The *DCS* of the C-term of *CB^clique^* is reduced and the N-term *CB^clique^* is very similar to the C-term one (**Figure 7c**). When considering all pathways, a single larger block (in red) is detected, containing 84% of the residues. The long-range pathway block is significantly larger than in the wild type, containing 46% of the residues from all the chains, extending toward the second half. Analysis of direct communications underlines a drastic increase of such communications between the helices at both halves, compared to the wild type (**Figure 7d,e**). Consequently, the E364F substitution significantly increases the stability of the communication network across the protein and results in a balance for the number of residues identified as *CB^clique^*.

## Conclusions

This study revealed that COMMA2 is able to detect a region known to be structurally ambiguous, *i.e*. well-ordered in a PDB structure and unresolved in another one, in the coiled-coil tetramer of the MeV PMD. The region is detected as a *CB^clique^* that spans all four chains. The application of COMMA2 to the coiled-coil tetramer of the NiV PMD yielded similar results, unveiling that the C-terminus part of the NiV PMD coiled-coil tetramer also has substantial disorder content. This property is not shared by the RhcC protein, which forms a right-handed tetrameric coiled-coil. Altogether, these results show that COMMA2 can be successfully applied to different protein systems to pinpoint specific and unique structural properties. Furthermore, comparisons with existing tools to predict disordered residues, revealed the power of COMMA2 to predict disordered residues, as well as to distinguish different dynamical behaviours (disorder, flexibility and stability) across the structure. Finally, this analysis enabled us to formulate a set of rules for the control of MeV and NiV PMD structural stability and inter-residue ‘communication’. Replacement of hydrophobic *a* positions with charged residues before the kink, leads to an increase in the number of direct communications in the second half and establishing communication between the two halves, which in turn enables the two halves to communicate across the structure. On the other hand, replacement of hydrophobic *a* positions with charged residues after the kink, leads to fragmentation of *CBs^path^* and decrease of communication in the second half and concomitant significant increase of communication in the first half. Learning from these rules, we proposed two contrasting substitutions. One at the kink-region, resulting in the overall increase of the flexibility through disruption of the communications. And a reverse substitution from a charged amino acid to a hydrophobic one at the C-term, leading to an increase in symmetry and communications between the two halves. Our results complement recent experiments on a set of substitutions of hydrophobic *a* or *d* positions resulting in non-functional proteins [11], by providing detailed descriptions of their dynamics. Furthermore, COMMA2 could help in understanding why some natural coiled-coils contain charged amino acids in *a* or *d* positions. COMMA2 provides a better understanding of proteins dynamics and is able to characterize disordered regions across protein structures. These results advocate for the application of COMMA2 to other protein families to highlight the role of disordered, ordered and flexible regions in their dynamics, and to determine the symmetrical or independent behaviour of the components.

## Author’s contributions

YK, EL and AC conceived the overall study and designed the experiments. YK implemented COMMA2. YK, PS and RV performed computational analysis. YK, EL and AC analysed the results and wrote the manuscript. DG and SL interpreted data, contributed to the manuscript preparation and approved the final draft. All of the authors have read and approved the final manuscript.

## Acknowledgements

This work was partially undertaken in the framework of CALSIMLAB supported by the public grant ANR-11-LABX-0037-01 constituting a part of the “Investissements d’Avenir” program (reference: ANR-11-IDEX-0004-02). It was also partially undertaken under the MAPPING project (ANR-11-BINF-0003, Excellence Programme “Investissement d’Avenir” in Bioinformatics). We acknowledge the access to the HPC resources of the Institute for Scientific Computing and Simulation at UPMC (Equip@Meso project - ANR-10-EQPX-29-01, Excellence Program “Investissement d’Avenir”).

