## Supporting Materials for "Predicting substitutions to modulate disorder and stability in coiled-coils"

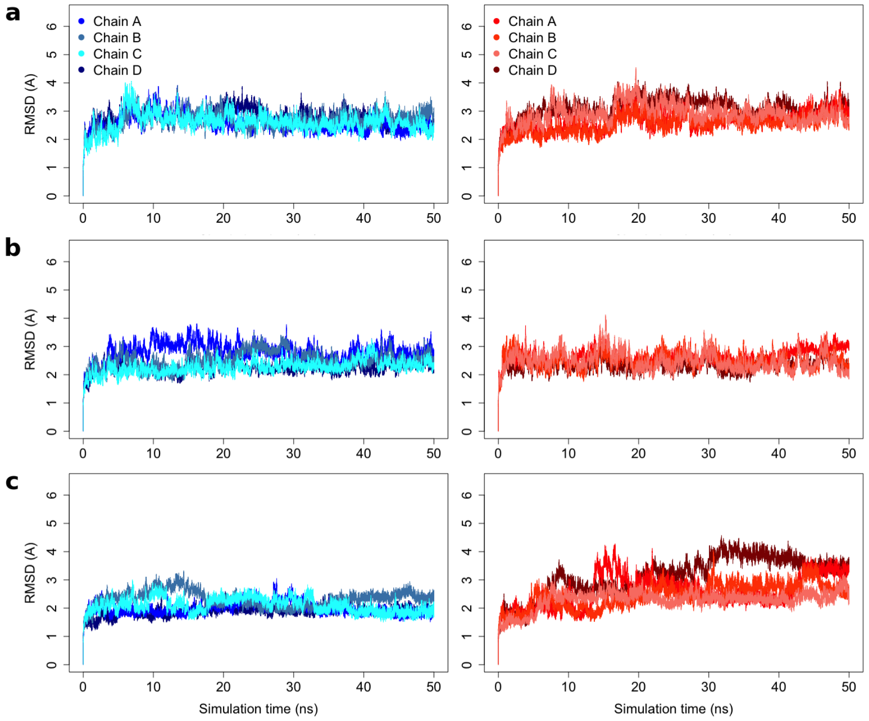


**Figure S1 RMSD plot for the studied coiled-coils.** The root mean square deviation is measured for every chain along the MD simulation time for the two replicates of **a)** MeV PMD (3ZDO), **b)** NiV PMD (4N5B), and **c)** RhcC (1YKB).

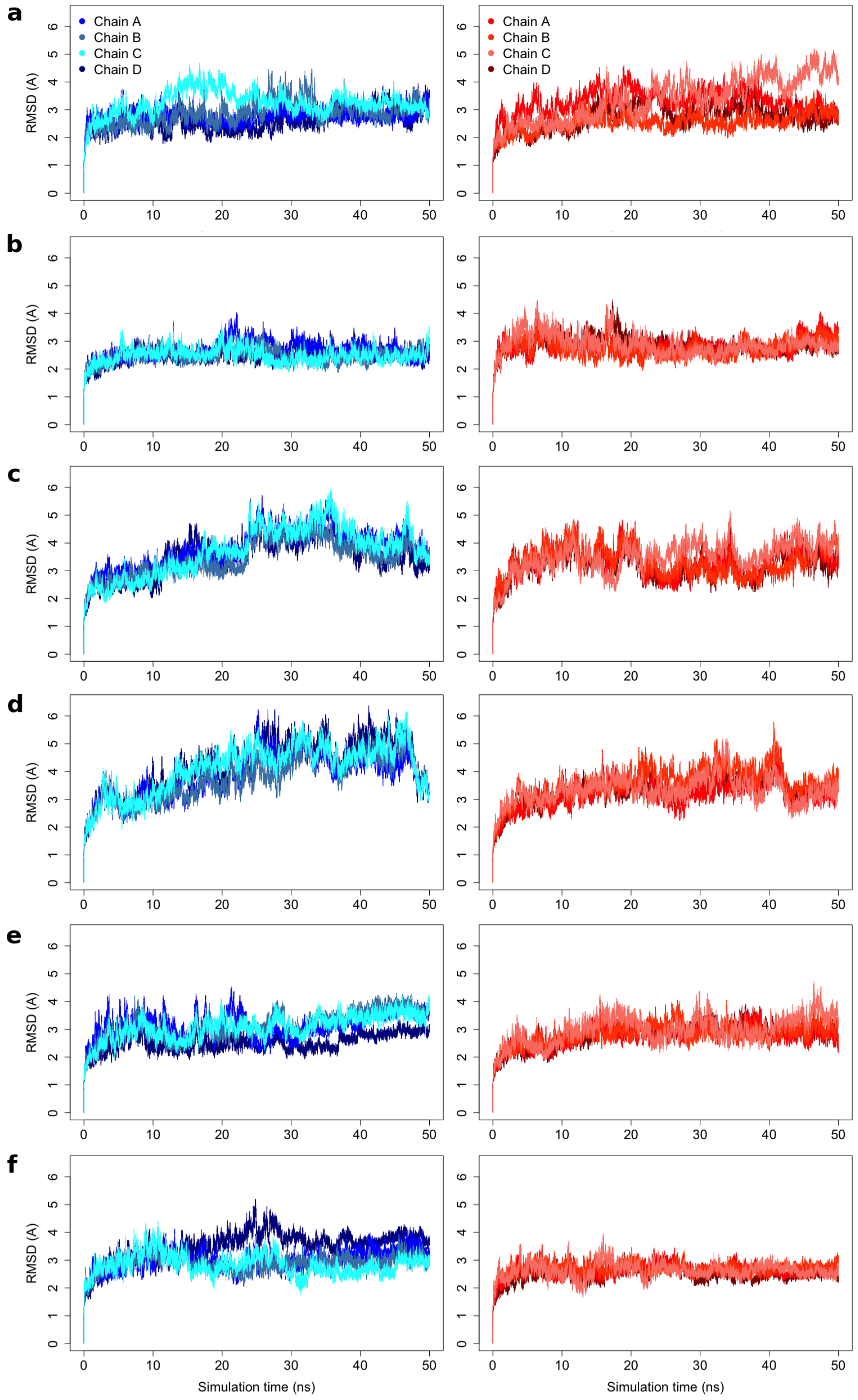


**Figure S2 RMSD plot for the variants of MeV PMD.** The root mean square deviation is measured for every residue along the MD simulation time for the variants of MeV PMD: **a)** V315D, **b)** L322D, **c)** L336D, **d)** V346D, **e)** I351D, and **f)** E364F.

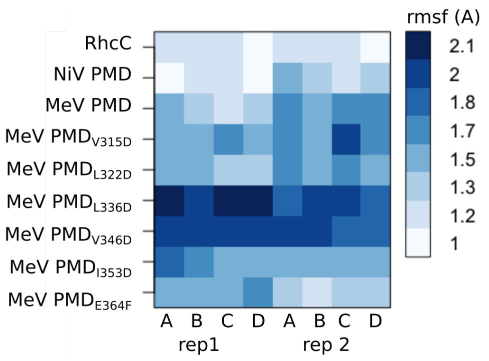


**Figure S3 RMSF plot for the studied coiled-coils.** The root mean square fluctuations is measured for every chain along the MD simulation time for the two replicates of NiV PMD (4N5B), RhcC (1YBK), MeV PMD (3ZDO) and six variants of MeV PMD.

### Comparison with single chains (for monomers)

One replicate of 50 ns MD trajectories was produced for each of the MeV PMD, NiV PMD and RhcC monomers, extracted from the tetramer crystal structures. The stability of the "artificial" systems were reached after 25 ns, therefore the last 25 ns of each replicate were chosen for analysis using COMMA2. In the simulations of both MeV and NiV PMDs, the unfolding of the C-term was observed. The single helices strongly bent during the simulation with the kink serving as a hinge point. The average structure obtained from MD simulations reveals the appearance of such a strong kink with high degree of bending for all the three systems (**Figure S4**). Several $CBs^{path}$ were observed for all the three monomers. Residues from 360 to 370, known to be structurally ambiguous in MeV PMD, were completely unfolded and detected as a $CB^{clique}$ by COMMA2 (**Figure S4d**, in blue). These results indicate that the C-terminal end of MeV and NiV PMD are intrinsically disordered. By contrast, in the case of RhcC monomer the C-term is folded, with no hints suggesting a propensity to disorder. We can interpret the results as a transition from unfolded state to "not-so-folded" state upon binding for the two PMDs.

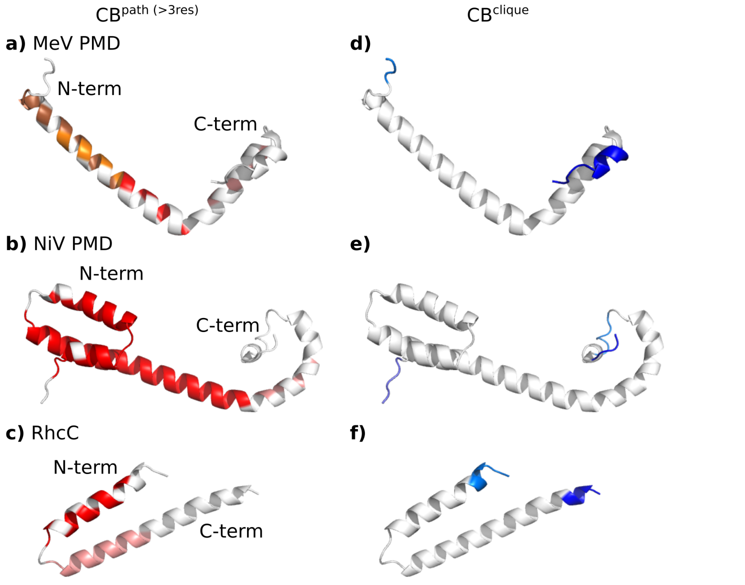


**Figure S4 Communication blocks identified by COMMA2 from single chain simulations.** Communication blocks are defined from applying COMMA2 to single chains of (**a,d**) MeV PMD, (**b,e**) NiV PMD and (**c.f**) RhcC. $CBs^{path}$ (**a,b,c**) identified by COMMA2 are mapped on the average MD conformation. Those pathways are obtained by considering interactions involving side chains and short-range pathways. The $CBs^{clique}$ (**d,e,f**) identified by COMMA2 are colored in blue tones.
